# Paired single-nucleus and spatial transcriptomics of the lung adenocarcinoma invasion sequence reveal niche domains and candidate reversal compounds

**DOI:** 10.64898/2026.10.08.757757

**Authors:** Zhiyang Li, Jiaxuan Yang

## Abstract

Lung adenocarcinoma (LUAD) develops through a defined morphological sequence from atypical adenomatous hyperplasia (AAH) to invasive adenocarcinoma, but the transcriptional programmes accompanying this transition have mostly been characterised in bulk tissue or in tumour tissue alone. We profiled paired samples spanning the full sequence by single-nucleus RNA sequencing (413,697 nuclei, 23 patients) and spatial transcriptomics (56 Visium sections, 25 patients), asking how the transcriptional state of the tissue changes as invasion proceeds and whether that state can be reversed *in silico* by known perturbations.

The atlas resolves into 39 second-level subtypes and the tissue into seven recurring niche-domain archetypes (*K*^∗^ = 7), six of which are ordinary lung structures and only one of which is restricted to invasive disease. A confound then shaped the remaining analyses: domain identity, marker stability and the apparent stage-dependence of the spatial programmes all scale with sequencing depth, and depth is in these sections a proxy for tissue density rather than a purely technical artefact (*ρ* = −0.89 between a domain’s depth and the fraction of its markers surviving depth matching). Several apparently positive results disappeared once depth was matched. The invasive domain’s ECM/interstitial identity survives that check: within patients the fibroblast ECM programme is higher in invasive disease in 20 of 22 paired cases (*P* = 0.0033 against random gene sets), and the result is unchanged when the genes added by hand are removed and under every leave-one-out variant.

Querying a patient-paired disease axis against the LINCS compound library with three scoring schemes returned 34 compounds in common, dominated by PI3K/mTOR inhibitors, with a split-half reproducibility of *ρ* = 0.962 against a permutation null of *±* 0.045 and an agreement of *ρ* = 0.735 between a single-cell-derived and a spatially derived axis. A positive control, the recovery of known regulators of an official benchmark (TBXT ranked 1st of 8,450), establishes that the pipeline recovers true positives, so the negative results reported here are properties of the data rather than of the method. Those negatives are stated explicitly: single-cell copy-number inference produced no usable malignant/non-malignant separator, the cohort-level spatial copy-number difference did not survive depth matching, a developmental trajectory ran opposite to that of the source study, gene-level prediction returned no signal on the pooled axis and only one perturbation survives compartment-wise gating, and a ligand-level analysis produced no usable pair.

## 1 Introduction

Lung adenocarcinoma (LUAD) develops through a stereotypic morphological sequence that begins with atypical adenomatous hyperplasia (AAH) and proceeds through adenocarcinoma *in situ* (AIS) and minimally invasive adenocarcinoma (MIA) to invasive adenocarcinoma (IAC). Each stage is histologically defined, and the transition from precursor lesion to invasive carcinoma is the point at which clinical management changes. The sequence therefore provides a well-defined system in which to ask what transcriptional changes accompany invasion and in what order they arise.

Most transcriptomic descriptions of this sequence derive either from bulk tissue (1) or from single-cell profiling restricted to the invasive tumour and adjacent normal lung, and both designs conflate the intermediate states that define the sequence. Single-cell atlases spanning the full spectrum are available (2), but these rarely carry a spatially resolved counterpart, so the tissue architecture surrounding a given transcriptional state, such as the niche composition around an alveolar type II cell, remains unobserved.

Two obstacles are specific to this disease. Tumour cells are difficult to identify without copy-number information, because the reference panels used for spatial deconvolution contain no malignant cell type; malignant epithelium is assigned to the most similar normal epithelial class, leaving any composition-based description of a niche structurally blind to the tumour. In addition, invasive regions of a section are both more cellular and more transcriptionally active than precursor regions, so comparisons between the two are also comparisons of RNA content.

We assembled paired single-nucleus RNA sequencing and Visium spatial transcriptomics from 23 patients spanning all five stages. The single-cell data define cell states and a patient-internal disease axis. The spatial data define tissue niches (3, 4) and provide the context in which the copy-number signal is placed (5). The same disease axis is then used as a query against two perturbation libraries (6), to ask whether any known compound or gene perturbation reverses it.

A methodological result obtained early shaped the remainder of the study. Sequencing depth and tissue density are not separable in this cohort, and their coupling is strong enough to generate stage-dependent patterns where none exist. Controlling for it, and reporting what survives when that control is incomplete, became the organising constraint of the work. We therefore present positive and negative results together and record the analytical difficulties encountered, several of which required the withdrawal of an apparently sound result.

## 2 Results

### 2.1 A single-nucleus atlas of the LUAD invasion sequence

Paired single-nucleus RNA sequencing and Visium spatial transcriptomics were obtained from 23 patients spanning the five stages of the sequence, with the spatial arm covering 56 sections from 25 patients and sharing 23 patients with the single-cell arm (Fig. 1a). Quality control retained 413,697 nuclei; after removal of contaminants and lowquality clusters, the six mutually exclusive lineage subsets on which the atlas is built contain 399,579 nuclei. The atlas resolves into six lineages (epithelial, fibroblast, endothelial, myeloid, T/NK, B/plasma; Fig. 1a) and, on per-lineage sub-clustering, into 39 second-level (L2) subtypes (Supplementary Table S1; Fig. 1f).

**Figure 1:**
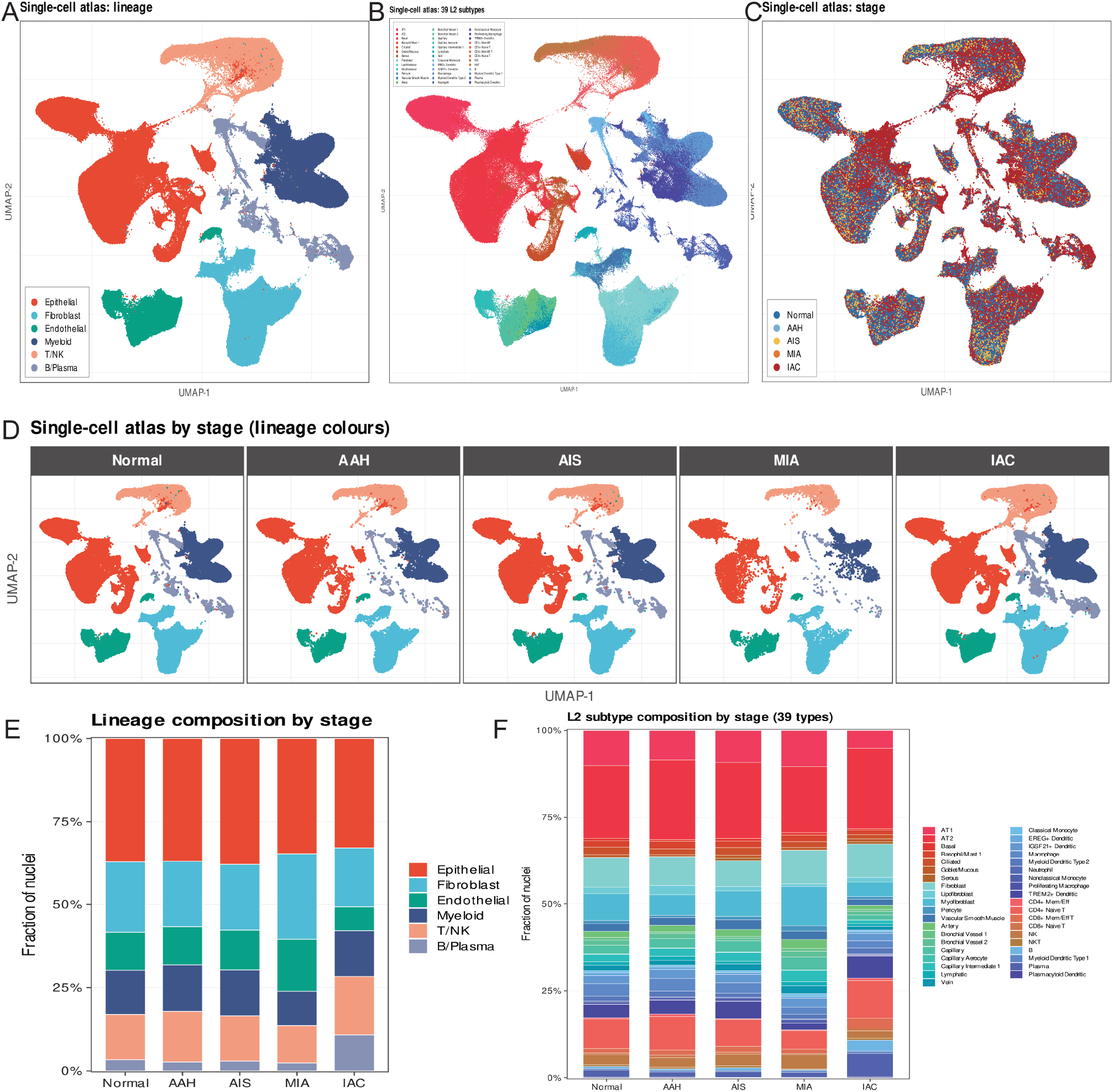
A single-nucleus atlas of the LUAD invasion sequence. (**a**) UMAP of 399,579 nuclei coloured by lineage. (**b**) Lineage composition across the cohort. (**c**) The same embedding coloured by disease stage. (**d**) The embedding split by stage, retaining lineage colours. (**e**) Lineage composition per stage. (**f**) Composition of the 39 second-level subtypes per stage. Endothelial and myeloid marker panels were rebuilt before composition analysis (Methods). The 39 subtype names are not unique lineage–subtype identifiers (67 combinations); compositions are reported at the lineage level in (**e**).

Two properties of the reference annotation affected the analyses that follow and are recorded here.

*Endothelial and myeloid panels were rebuilt*. The published marker table supplies six canonical genes per endothelial subtype, which is insufficient to discriminate between closely related endothelial states. Under that panel the *Artery* subtype was assigned to 16 clusters across the cohort; under an expanded panel of up to 20 genes per subtype across nine endothelial types it was assigned to four, with the remainder reassigned to *Lymphatic, Vein* and *Capillary*. The endothelial and myeloid panels were therefore rebuilt before any downstream analysis. Analyses that depend on endothelial subtype proportions inherit the corrected panels.

*The L2 nomenclature is not a set of unique identifiers*. The 39 names map onto 67 distinct lineage–subtype combinations. The clearest case is *Basophil/Mast 1*, which the reference assigns predominantly to epithelial cells (6,360 cells) and only marginally to myeloid cells (8 cells), so the name misrepresents the content. Subtype composition is accordingly reported at the lineage level (Fig. 1e).

A second difficulty concerns the coverage rather than the nomenclature of the reference. The 39-subtype reference contains no exhaustion programme, and exhausted T cells were consequently absorbed into the nearest available label. One cluster carries LAG3, TIGIT, HAVCR2, ENTPD1, GZMB and PRF1 while lacking CCR7, SELL, TCF7 and LEF1, an exhausted rather than a naive phenotype, yet the reference assigns it to *CD8+ Naive T*. Deconvolution cannot recover a state absent from the reference, so the T-cell composition reported here is a composition in the reference’s vocabulary. Functional exhaustion was therefore measured independently, by projecting an exhaustion signature (7, 8) onto the spots.

One AAH sample (P24) carried a plasma-cell programme confined to a single patient and not tracking disease stage. A patient-level rank of B/plasma content, comparison against a matched negative-control gene set, and an independent readout from a separate region of the same block each identified it as ambient RNA rather than a tumour-associated infiltrate. The sample was retained with an annotation, and all conclusions that depend on B/plasma content are reported both with and without it. The source study does not describe ambient-RNA correction, and P24 is one of its representative samples.

### 2.2 Spatial deconvolution resolves seven recurring niche domains

BANKSY applied per section yielded 434 spatial domains, which were collapsed to archetypes by clustering their expression profiles. The stability of that collapse was measured by subsampling: re-clustering on a random 80% of the domains, repeated 100 times, and comparing to the full solution. This gives 0.945 at *K*^*∗*^ = 7, the largest *K* above the 0.90 threshold, and 0.966 at *K* = 6; it falls to 0.814 at *K* = 8, so the archetypes cannot be subdivided further. We therefore treat *K*^*∗*^ = 7 as the resolution limit of this cohort. A purely descriptive optimum exists at *K* = 6 and was not adopted. These figures describe the archetype clustering and are *not* a cross-seed measurement; the cross-seed agreement of the underlying per-section domain calls is a separate and lower quantity (median adjusted Rand index 0.782), and is what determines the stability gate reported in the Limitations.

Identities were assigned from each archetype’s RCTD composition (Fig. 2a,c) and its marker genes (Fig. 2g). Six of the seven are ordinary lung structures: airway epithelium (D1), peribronchovascular interstitium carrying an inflammatory CAF programme (D2) (9, 10), alveolar wall (D4), AT2 epithelium (D5), vascular and airway smooth muscle (D6), and lymphocyte aggregates (D7). Only D3 is restricted to invasive disease. On the stage axis (Fig. 2d), D3 accounts for 17% of spots in IAC and **0% in every precursor stage**, whereas D7 is present in all five stages. The presence of lymphoid aggregates before invasion argues against the assumption that immune aggregation accompanies malignancy.

**Figure 2:**
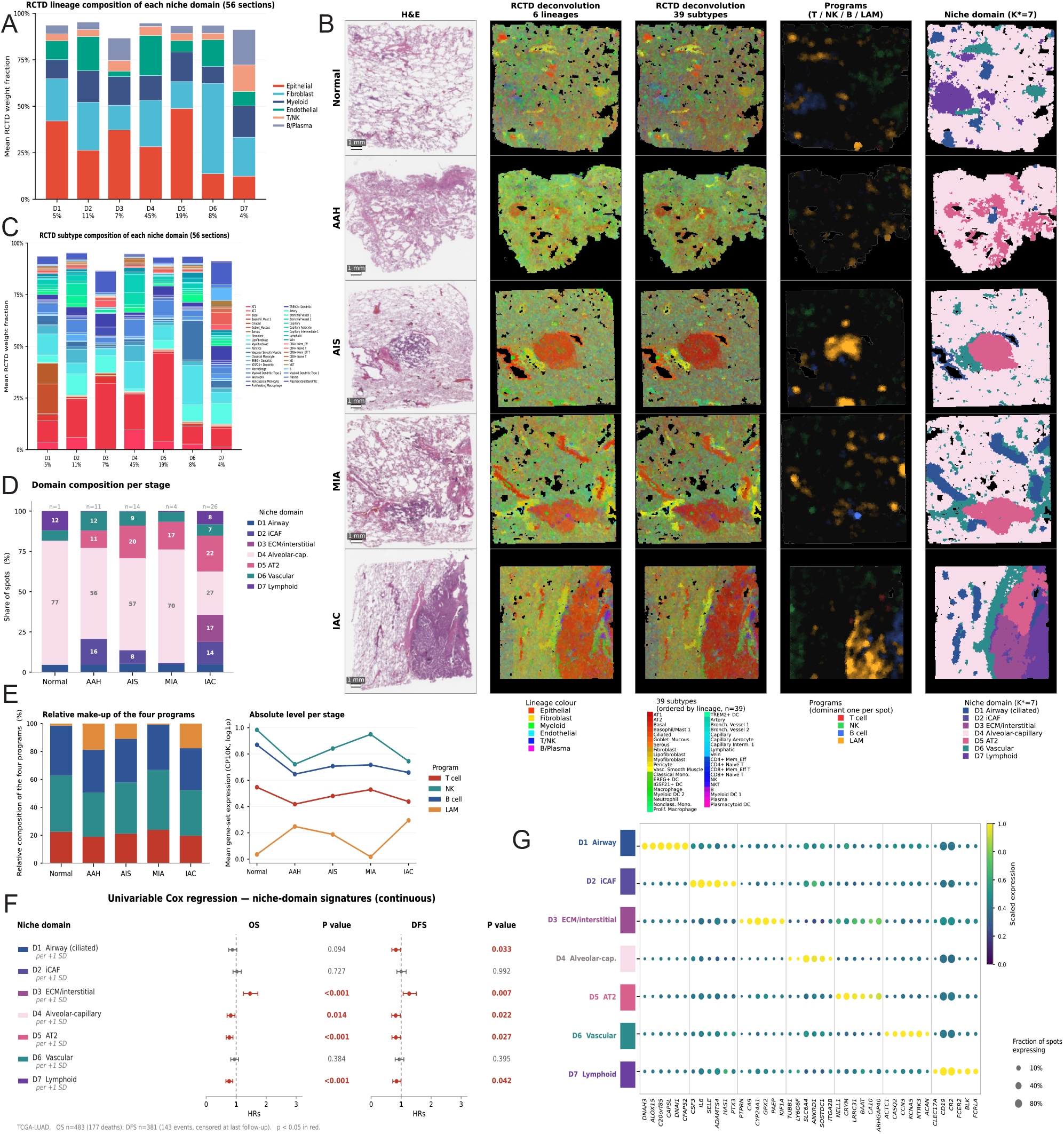
Spatial deconvolution resolves seven niche-domain archetypes. (**a**) RCTD lineage composition of each domain archetype (56 sections). (**b**) Spatial matrix: one representative section per stage (rows) against H&E, RCTD at the 6-lineage and 39-subtype level, the four spatial programmes (dominant programme per spot) and the niche-domain archetype (columns). Scale bars are calibrated per section from the nominal 100-*μ*m spot spacing. (**c**) RCTD subtype composition of each domain archetype. (**d**) Domain composition per stage; D3 is confined to invasive disease (17% of spots, 0% in all precursor stages) whereas the lymphoid domain D7 occurs in all five stages. (**e**) Relative make-up and absolute level of the four spatial programmes per stage. **This panel is a negative control**: before depth matching the programme scores essentially mirror sequencing depth, and after matching into depth deciles the stage ordering is identical across all ten deciles but does not follow disease progression. The fourth programme is the lipid-associated macrophage (LAM) signature (11), which we report alongside three exhaustion axes; it is not an exhaustion signature in the T-cell sense. (**f**) Univariable Cox regression of each domain’s marker signature on overall and disease-free survival (TCGA-LUAD); p *<* 0.05 in red. The D3 row uses the unadjusted signature (see main text). (**g**) Marker-gene dot plot for the seven domains.

Two caveats attach to the domain labels. The published marker table for these domains is itself depthconfounded (see below), and the identity of D3 was revised twice during the study, from an immune-infiltrate reading to a hypoxic/invasive reading and then to its current ECM/interstitial reading; the earlier names should not be used. The domain definition is also method-sensitive: BANKSY, RCTD composition and marker-based clustering return 7, 6 and 5 domains respectively, and the expression-based partition agrees with an RCTD-aligned partition at an adjusted Rand index of 0.352. Domain boundaries are therefore quantitative but not unique. The composition underlying each label is given in Table 1.

**Table 1:** Identity of the seven niche-domain archetypes. Enrichment is the RCTD composition of a domain relative to the rest of the cohort, as a fold change; the share is the fraction of all spots assigned to that domain. The D3 label was revised twice during the study (see main text) and the earlier names should not be used. No label here implies a malignancy call.

| Domain | Label | Basis (RCTD enrichment or depth-balanced markers) | Share | Note |
| --- | --- | --- | --- | --- |
| D1 | Airway epithelium | Ciliated $\times 6.27$; Goblet $\times 5.60$ | 5% | |
| D2 | iCAF stroma | IL6, CXCL8, CCL2, LIF, HAS1, HAS2 | 11% | *peribronchovascular* |
| D3 | ECM / interstitial | COL1A1, COL3A1, FN1, CTHRC1 | 17% (IAC) | *no precursor stage; two label revisions* |
| D4 | Alveolar wall | Capillary $\times 3.56$; AT1 $\times 2.43$ | 45% | *77% of the Normal section* |
| D5 | AT2 epithelium | AT2 weight 0.426 ($\times 2.16$) | 19% | *highest copy-number score* |
| D6 | Vascular / airway wall | VSMC $\times 4.26$; Artery $\times 2.33$ | 8% | |
| D7 | Lymphoid aggregate | B $\times 4.82$; NKT $\times 3.13$ | 4% | *present in all five stages* |

### 2.3 Sequencing depth is a proxy for tissue density

The seven domains differ in sequencing depth by a factor of 1.00 to 7.95. Depth matching removes a large and predictable fraction of each domain’s marker genes: across the seven domains, the fraction of a domain’s top-150 upgenes surviving per-slide depth matching falls monotonically with depth (Spearman *ρ* = *−* 0.89; Fig. 3f), with two independent estimators (decile matching and per-gene regression on log_10_(nUMI)) agreeing almost gene-for-gene. D3 is the sole outlier; it is both the deepest domain and the only one restricted to invasive disease, and 16 of its 150 markers survive matching.

**Figure 3:**
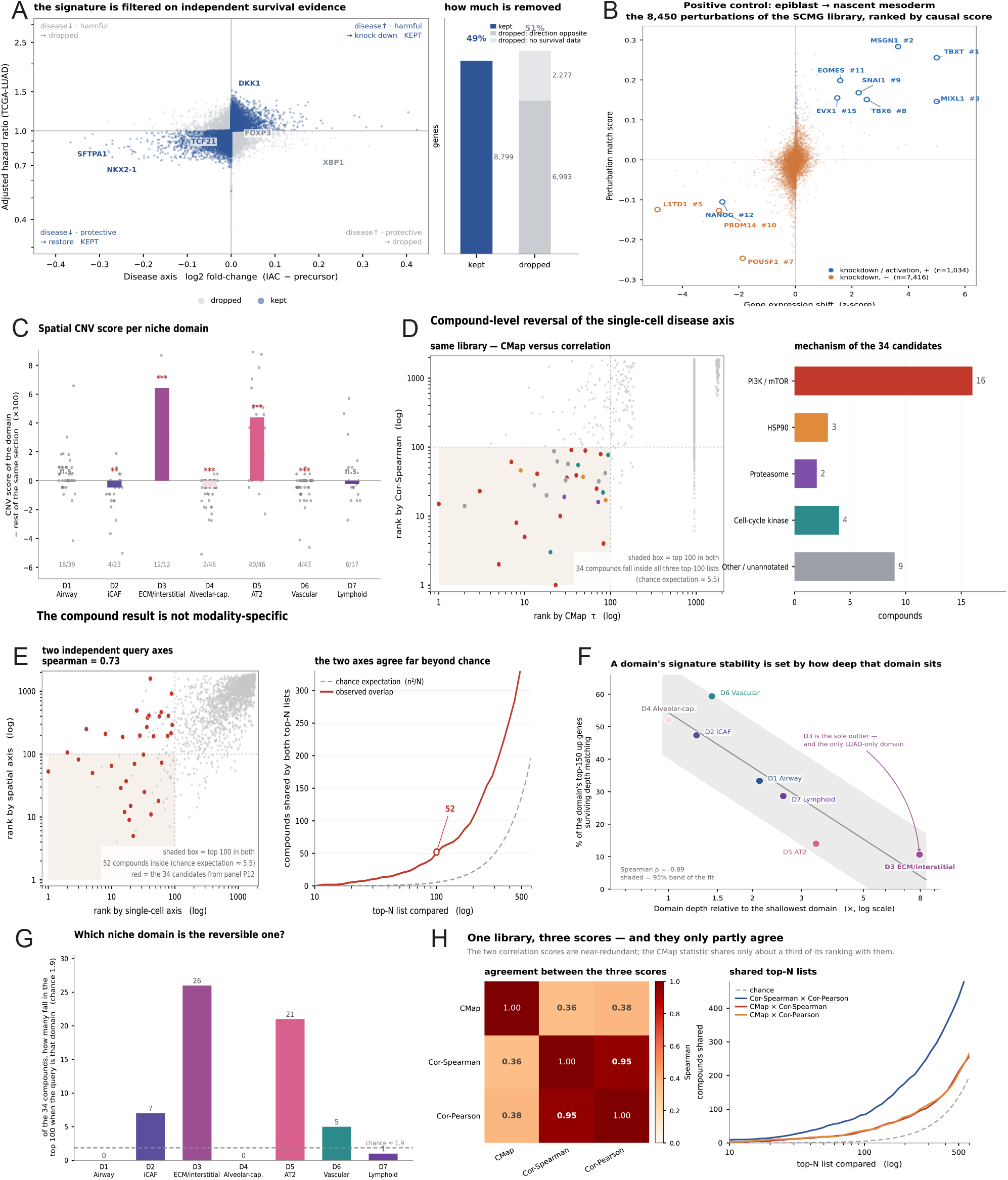
*In-silico* reversal of the disease state. (**a**) The query signature is filtered on independent survival evidence: disease direction against adjusted hazard ratio, with genes whose directions disagree set to zero. (**b**) Positive control — the official epiblast-to-nascent-mesoderm benchmark, with the known regulators labelled and ranked (TBXT 1st of 8,450). (**c**) Spatial copy-number score per domain, each domain compared with the remaining spots of the same section so that all spots in a comparison share one library. (**d**) Compound-level reversal: CMap rank against Cor-Spearman rank for the 1,826 compounds, with the 34 compounds common to all three scoring schemes highlighted, and their mechanism composition. (**e**) The candidate ranking is not modality-specific: a single-cell-derived and a spatially derived query axis agree at Spearman *ρ* = 0.735. (**f**) A domain’s marker stability scales with its sequencing depth (*ρ* = *−* 0.89; D3 is the sole outlier and the only invasive-restricted domain). (**g**) Of the 34 candidates, 26 fall in the top 100 for the ECM/interstitial domain and 21 for the AT2 domain, against a chance expectation of 1.9. (**h**) One library, three scores: the two correlation scores are near-redundant (*ρ* = 0.950) whereas the CMap statistic shares only about a third of its ranking with them.

This resembles a standard technical confound, and treating it as one would be an error. The same quantity is independently visible in the images: the PLIP-derived neoplastic score correlates with the fraction of a spot occupied by tissue at *ρ* = 0.754 across four sections of one patient.

Deep spots are therefore deep because they are dense, and dense because they contain tumour. Removing depth as a covariate removes part of the biological signal; the operation is closer to over-adjustment than to deconfounding, and neither the unadjusted nor the adjusted estimate is correct on its own.

This coupling is the binding constraint of the study and recurs in four places. In the spatial copy-number arm it determines which comparisons survive. In the prognosis arm it is the reason one domain’s apparent prognostic effect shrinks when its signature is replaced by a depth-balanced version. In the developmental analysis it accounts for a trajectory running opposite to that of the source study. It also prevents interpretation of the stage-dependence of the four spatial programmes. Before matching, their scores are almost a mirror of depth (per-spot *r* =*−* 0.30 to *−* 0.69 for the T-cell programme; sections differ 2.4-fold in median depth). After matching into depth deciles the stage ordering is identical across all ten deciles but does not follow disease progression: AAH and AIS are both precursor stages yet occupy opposite ends of the ordering. That comparison is consequently reported as a negative control (Fig. 2e).

Because the revised identity of D3 is an ECM/interstitial programme, that programme was tested directly rather than accepted from the domain analysis. Within each patient, fibroblasts were scored against the cohort fibroblast mean and standard deviation per gene, and the mean of a 22-gene ECM set was compared between invasive and precursor fibroblasts. The programme is higher in invasive disease in **20 of the 22 patients with both stages** (median within-patient difference +0.402; 300 random gene sets of matched size drawn from expressed genes give *P* = 0.0033 on both the count and the median). Eight of the 22 genes were added by us rather than drawn from the depth-balanced D3 ranking; restricting the panel to the 14 data-derived genes gives 20 of 22 (+0.347, *P* = 0.0033), restricting it to the eight hand-added genes also gives 20 of 22 (+0.450, *P* = 0.0033), and removing any single gene leaves the count at 20 of 22 in all 22 leave-one-out variants. The ECM programme is therefore not an artefact of gene selection, although the result describes a tissue-level programme in fibroblasts and does not identify which ECM component is responsible.

### 2.4 Copy-number inference: three failures and one surviving signal

Malignant-cell identification is where the analytical failures of this study concentrate, and they are reported in full because they constrain the interpretation of the spatial results.

#### Single-cell inference

Across six tool families, comprising CopyKAT (12) in default and floored configurations, two anchor-based arms, inferCNV v1 and v2 (13), and a supervised expression-based classifier (14), no configuration produced a usable malignant/non-malignant separator, and each failure was traced to a specific cause. The clearest was an instability check: the same set of 3,890 normal cells, used as the reference in three separate runs, produced aneuploid calls of 0%, 13.6% and 49.8%. The most misleading was an AUROC of exactly 0.5000 from inferCNV v1, where all 19,748 cells carried a single identical score value because the per-cell statistic reduced to the constant reference baseline; the ROC was a degeneracy rather than evidence of an absent difference. A second version produced AUROC 0.652 but failed its own control, the normal cells scoring 0.0826 against a threshold of 0.05.

#### Cohort-level spatial copy number

All 25 cases ran successfully at the Visium level. The cohort-level claim that invasive sections carry more copy-number signal shrank by 71% after depth balancing, and the matched comparison was not significant (*p* = 0.092, confidence interval including zero). A random subsample reproduced the pre-matching result index by index, confirming that the original effect was driven by sequencing depth rather than by sample size. No cohort-level copy-number difference is therefore reported.

#### One comparison survives by design

Comparing each domain against the remaining spots of the same section removes depth by construction, since every spot in the comparison shares one cDNA library (Fig. 3c). Under this design the copy-number score is elevated in D5 (AT2; median Δ = +0.044, positive in 40 of 46 sections, *p <* 0.001) and in D3 (12 of 12 sections, *p* = 0.0005); D4, D6 and D2 are significantly lower, and D1 and D7 show no effect. Restricted to the 25 precursor sections, D5 remains positive in 21, so the signal is present before invasion. This is consistent with AT2 as the cell of origin in this disease and with an independent supervised classifier whose signal was likewise confined to AT2.

The score is a continuous quantity and not a malignancy call; no malignant label is assigned to any spot or domain.

Within D5, splitting spots into high and low copy-number groups produces spatially coherent patches, but the split is not separable from RNA content: the correlation between score and total UMI within D5 is *r* = 0.755 across sections, and splitting by spot counts alone reproduces roughly 75% of the difference in AT2 content between the two groups. A “D5-high” subdomain therefore cannot be described as a tumour subdomain.

### 2.5 Prognostic association of the niche domains

The domain marker signatures were tested for prognostic information in an independent cohort (TCGA-LUAD; 483 patients, 177 deaths; 381 patients and 143 events for disease-free survival). Using each domain’s top-30 markers standardised across samples, the associations are consistent between a continuous parameterisation and a median split (Fig. 2f) and between overall and disease-free survival. A high D7 (lymphoid) signature is associated with better outcome (continuous HR 0.77, *p <* 0.001; median split HR 0.60) and a high D3 signature with worse (HR 1.46, *p <* 0.001; HR 2.03); D4 and D5 are protective on overall survival.

The D3 result requires qualification. Its signature is drawn from the raw marker table, which the depth analysis shows to be dominated by depth for this domain. Replacing it with the depth-balanced signature reduces the adjusted hazard ratio from 2.30 to 1.30 (*p* = 0.09), and the connectivity-map signal for the same signature falls within the random range. The direction and its robustness are therefore reported, but not the point estimate, and D3 is not treated as an established prognostic feature.

### 2.6 *In-silico* reversal of the disease state

To ask whether the disease state can be reversed, a query signature was derived from the single-cell data and used against two perturbation libraries.

#### Query construction

For each patient the difference in mean expression between invasive and precursor cells was computed and the median taken across patients, giving an 18,069-gene axis, of which 16,104 are represented in the SCMG library. This axis was filtered against an independent survival signal (per-gene adjusted Cox models in TCGA-LUAD), retaining a gene only where the direction of disease change agreed with the direction of prognostic effect, so that a gene rising in disease is retained only if higher expression also predicts worse outcome, and conversely (Fig. 3a). The filter removes 9,270 of 18,069 genes (51% of the full set; 44% of the 15,792 genes with survival data), of which 6,993 change in disease but are protective and 2,277 lack survival annotation. It is applied because a reversal query that ignores it would target protective programmes as readily as harmful ones.

#### Scoring schemes

The signature was scored against the LINCS library using the CMap weighted-KS statistic (*τ* ) and two correlation-based scores. The library was rebuilt and held locally rather than queried through the CLUE scoring service, which was retired on 2026-01-31 (Methods), so the version queried is fixed and re-runnable and every scoring path is cross-validated against the official implementation. The two correlation scores are nearly redundant (Spearman *ρ* = 0.950; 433 of 1,826 compounds shared in their top 500), whereas the CMap statistic shares only about a third of its ranking with them (*ρ* = 0.36–0.38; 18–19 of 50 in the top 50) (Fig. 3h). This reflects the statistics rather than the implementation: CMap-type scores use only the membership of the query’s gene sets and discard the perturbation values, and are therefore systematically different from value-based correlations. Requiring a compound to rank in the top 100 under all three schemes is consequently a stricter filter than it appears, and yields **34 compounds** (chance expectation 5.5), of which 16 (47%) target PI3K/mTOR, with smaller groups against HSP90, the proteasome and cell-cycle kinases (Fig. 3d; Table 2).

**Table 2:** The 34 compounds common to all three scoring schemes. Compounds rank in the top 100 under each of the CMap weighted-KS statistic (*τ* ) and the two correlation scores, applied to the prognosis-filtered patient-internal disease axis against the LINCS library (1,826 compounds). Rows are ordered by CMap rank. Values are medians across the cell lines in which a compound was profiled; more negative indicates stronger reversal. The chance expectation for a compound to appear in all three top-100 lists is 5.5. Mechanisms are annotated from library metadata, curated for incompletely annotated compounds; the assignment plays no part in selection.

| Compound | Mechanism | CMap $\tau$ | Cor-Sp $\rho$ | Cor-Pr $\rho$ | Rank CMap | Rank Cor-Sp |
| --- | --- | --- | --- | --- | --- | --- |
| AZD-2014 | PI3K / mTOR | -0.355 | -0.186 | -0.112 | 1 | 15 |
| lacidipine | PI3K / mTOR | -0.330 | -0.188 | -0.113 | 2 | 14 |
| AZD-8055 | PI3K / mTOR | -0.329 | -0.176 | -0.111 | 3 | 23 |
| PKI-179 | PI3K / mTOR | -0.322 | -0.223 | -0.152 | 5 | 2 |
| GDC-0941 | PI3K / mTOR | -0.311 | -0.111 | -0.071 | 7 | 61 |
| MLN-0128 | PI3K / mTOR | -0.311 | -0.205 | -0.128 | 8 | 8 |
| NVP-AUY922 | HSP90 | -0.304 | -0.135 | -0.091 | 9 | 46 |
| GDC-0980 | PI3K / mTOR | -0.302 | -0.215 | -0.137 | 10 | 5 |
| BIIB-021 | JAK2 | -0.294 | -0.170 | -0.118 | 13 | 28 |
| taselisib | PI3K / mTOR | -0.291 | -0.146 | -0.116 | 14 | 41 |
| okadaic acid | phosphatase | -0.286 | -0.179 | -0.142 | 18 | 20 |
| XL-888 | cell-cycle kinase | -0.280 | -0.223 | -0.134 | 20 | 3 |
| JW-7-24-1 | mTOR | -0.279 | -0.094 | -0.065 | 22 | 87 |
| torin-2 | PI3K / mTOR | -0.277 | -0.237 | -0.153 | 23 | 1 |
| midostaurin | kinase | -0.276 | -0.109 | -0.059 | 24 | 62 |
| NVP-BEZ235 | PI3K / mTOR | -0.274 | -0.199 | -0.124 | 26 | 10 |
| bortezomib | proteasome | -0.271 | -0.180 | -0.106 | 29 | 19 |
| pentobarbital | GABA-A | -0.266 | -0.163 | -0.093 | 30 | 33 |
| PI-103 | PI3K / mTOR | -0.264 | -0.155 | -0.093 | 31 | 36 |
| TAK-285 | HER2 / EGFR | -0.261 | -0.117 | -0.081 | 32 | 57 |
| voxtalisib | PI3K / mTOR | -0.257 | -0.093 | -0.063 | 35 | 91 |
| torin-1 | PI3K / mTOR | -0.250 | -0.149 | -0.094 | 40 | 39 |
| PF-03814735 | Aurora kinase | -0.242 | -0.119 | -0.067 | 42 | 55 |
| AT-13387 | HSP90 | -0.237 | -0.154 | -0.089 | 48 | 37 |
| OSI-027 | mTOR | -0.236 | -0.093 | -0.062 | 51 | 89 |
| WYE-125132 | mTOR | -0.217 | -0.172 | -0.107 | 69 | 25 |
| MG-132 | proteasome | -0.214 | -0.184 | -0.109 | 72 | 16 |
| NVP-TAE226 | FAK | -0.213 | -0.164 | -0.101 | 73 | 32 |
| GSK-2110183 | PI3K / mTOR | -0.209 | -0.101 | -0.081 | 77 | 79 |
| dinaciclib | CDK | -0.208 | -0.177 | -0.102 | 82 | 22 |
| GSK-2126458 | PI3K / mTOR | -0.208 | -0.218 | -0.139 | 83 | 4 |
| YM-155 | survivin | -0.205 | -0.140 | -0.075 | 87 | 42 |
| tanespimycin | HSP90 | -0.205 | -0.183 | -0.118 | 88 | 17 |
| SB-939 | cell-cycle kinase | -0.199 | -0.102 | -0.059 | 94 | 77 |

**Table 3:**
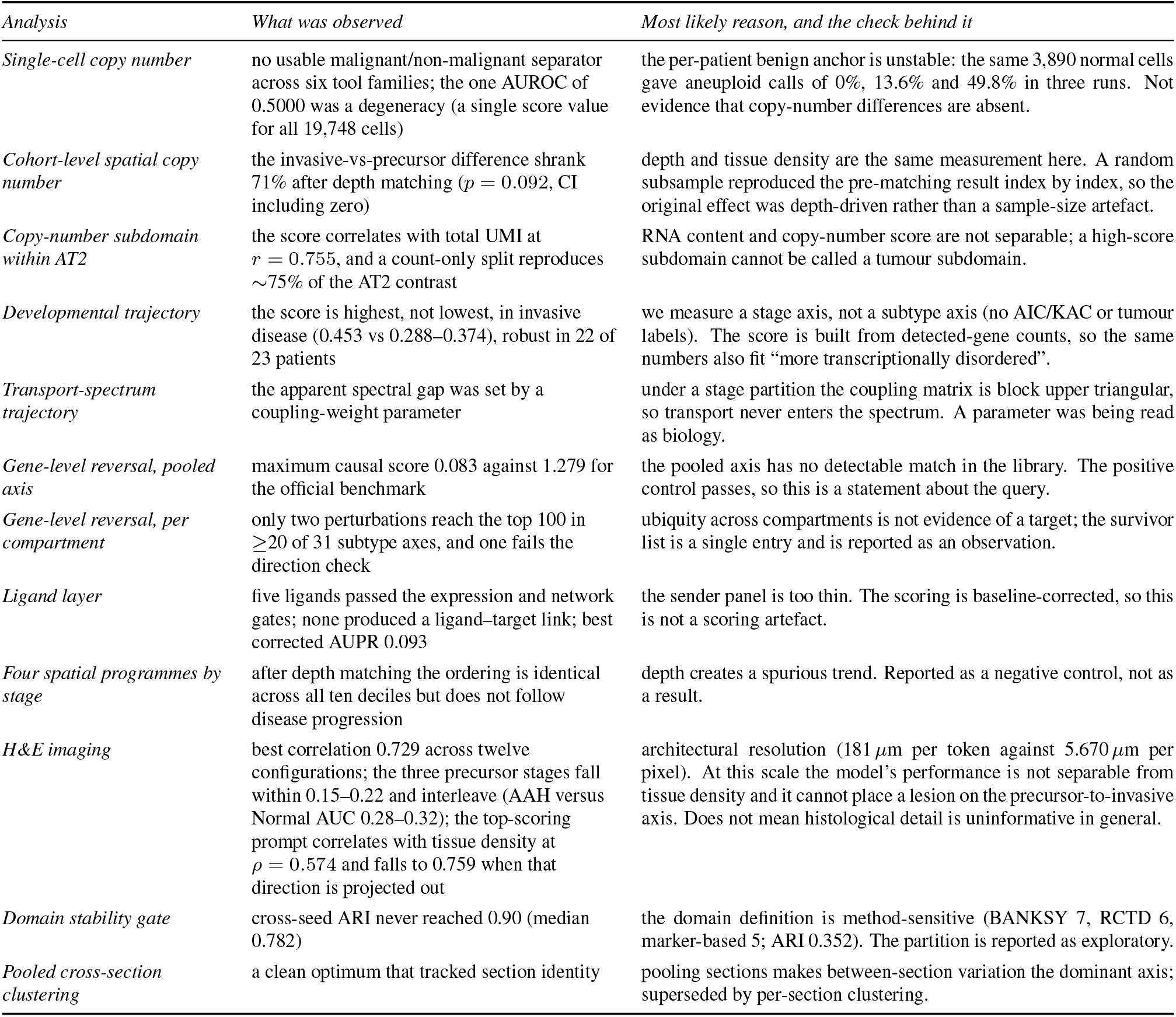
Negative results and the reason we believe each one failed. The right-hand column is an assessment, not a measurement: it names the most likely explanation and, where we could test it, the check that separates an absent signal from a broken method. The distinction matters because a negative result with a passing positive control constrains the biology, whereas one without does not.

| *Analysis* | *What was observed* | *Most likely reason, and the check behind it* |
| --- | --- | --- |
| *Single-cell copy number* | no usable malignant/non-malignant separator across six tool families; the one AUROC of 0.5000 was a degeneracy (a single score value for all 19,748 cells) | the per-patient benign anchor is unstable: the same 3,890 normal cells gave aneuploid calls of 0%, 13.6% and 49.8% in three runs. Not evidence that copy-number differences are absent. |
| *Cohort-level spatial copy number* | the invasive-vs-precursor difference shrank 71% after depth matching ($p = 0.092$, CI including zero) | depth and tissue density are the same measurement here. A random subsample reproduced the pre-matching result index by index, so the original effect was depth-driven rather than a sample-size artefact. |
| *Copy-number subdomain within AT2* | the score correlates with total UMI at $r = 0.755$, and a count-only split reproduces $\sim 75\%$ of the AT2 contrast | RNA content and copy-number score are not separable; a high-score subdomain cannot be called a tumour subdomain. |
| *Developmental trajectory* | the score is highest, not lowest, in invasive disease (0.453 vs 0.288–0.374), robust in 22 of 23 patients | we measure a stage axis, not a subtype axis (no AIC/KAC or tumour labels). The score is built from detected-gene counts, so the same numbers also fit “more transcriptionally disordered”. |
| *Transport-spectrum trajectory* | the apparent spectral gap was set by a coupling-weight parameter | under a stage partition the coupling matrix is block upper triangular, so transport never enters the spectrum. A parameter was being read as biology. |
| *Gene-level reversal, pooled axis* | maximum causal score 0.083 against 1.279 for the official benchmark | the pooled axis has no detectable match in the library. The positive control passes, so this is a statement about the query. |
| *Gene-level reversal, per compartment* | only two perturbations reach the top 100 in $\geq 20$ of 31 subtype axes, and one fails the direction check | ubiquity across compartments is not evidence of a target; the survivor list is a single entry and is reported as an observation. |
| *Ligand layer* | five ligands passed the expression and network gates; none produced a ligand–target link; best corrected AUPR 0.093 | the sender panel is too thin. The scoring is baseline-corrected, so this is not a scoring artefact. |
| *Four spatial programmes by stage* | after depth matching the ordering is identical across all ten deciles but does not follow disease progression | depth creates a spurious trend. Reported as a negative control, not as a result. |
| *H&E imaging* | best correlation 0.729 across twelve configurations; the three precursor stages fall within 0.15–0.22 and interleave (AAH versus Normal AUC 0.28–0.32); the top-scoring prompt correlates with tissue density at $\rho = 0.574$ and falls to 0.759 when that direction is projected out | architectural resolution ($181\ \mu\text{m}$ per token against $5.670\ \mu\text{m}$ per pixel). At this scale the model’s performance is not separable from tissue density and it cannot place a lesion on the precursor-to-invasive axis. Does not mean histological detail is uninformative in general. |
| *Domain stability gate* | cross-seed ARI never reached 0.90 (median 0.782) | the domain definition is method-sensitive (BANKSY 7, RCTD 6, marker-based 5; ARI 0.352). The partition is reported as exploratory. |
| *Pooled cross-section clustering* | a clean optimum that tracked section identity | pooling sections makes between-section variation the dominant axis; superseded by per-section clustering. |

#### Reproducibility and modality independence

Splitting patients at random into halves, deriving the axis independently in each half and scoring both against LINCS reproduces the ranking with a median Spearman *ρ* of 0.962 across four splits, against a permutation null of *±* 0.045 (Fig. 3e, left); the top-500 compound lists share 423 on median against a chance expectation of 137. One split is weaker (*ρ* = 0.752) and all four are reported. A second query axis was derived from the spatial data, after depth balancing, by comparing invasive and precursor spots within nUMI deciles, and scored against the same library with the same statistic. The two axes agree at Spearman *ρ* = 0.735 (top-100 overlap 52, chance expectation 5.5) (Fig. 3e, right). Since the two axes derive from different platforms and different pairing strategies, this is a stronger reproducibility check than repeating the scoring scheme within one modality.

#### Positive control

Before the candidate list was examined, the pipeline was verified against an official benchmark with a known answer, the epiblast-to-nascent-mesoderm transition, using the authors’ own implementation, manifold and standard deviations (15). The known regulators are recovered at the top of the ranking: TBXT 1st of 8,450, MSGN1 2nd, MIXL1 3rd, POU5F1 7th, SNAI1 9th, EOMES 11th, NANOG 12th and EVX1 15^th^ (Fig. 3b). This establishes that an empty result from the pipeline indicates the query or the data rather than a broken method. Numerical identity with the official implementations was confirmed separately: the weighted-KS implementation reproduces the reference to a maximum absolute difference of 6.6 *×* 10^*−*14^ with 2000 of 2000 signs agreeing, and the vectorised causal score reproduces the authors’ Python at Spearman 1.00000000. Table 4 collects these and the other numerical agreements reported below.

**Table 4:** Method verification and positive controls. Every numerical check compares our own implementation with the reference implementation on identical input, so a discrepancy would be an implementation error rather than a modelling choice. The biological control establishes that an empty result from this pipeline reflects the query and not the method.

| *Check* | *Compared against* | *Result* |
| --- | --- | --- |
| *Numerical agreement with the reference implementation* |  |  |
| Weighted-KS statistic | signatureSearch 1.12.0 | max. abs. difference $6.600 \times 10^{-14}$; signs agree 2000.000/2000.000 |
| Normalised connectivity | signatureSearch 1.12.0 | max. abs. difference $1.200 \times 10^{-13}$ |
| Causal gene score | authors’ Python | Spearman 1.000; signs agree 100% (19,819.000 entries) |
| *Biological positive control* |  |  |
| Official benchmark | epiblast → nascent mesoderm | TBXT 1st of 8450.000; MSGN1 2nd; MIXL1 3rd; POU5F1 7th; SNAI1 9th; EOMES 11th; NANOG 12th; EVX1 15th |
| *Stability and agreement* |  |  |
| Split-half | 4 patient partitions | median $\rho = 0.962$ (one split 0.752); permutation null $\pm 0.045$ |
| Cross-modality | single-cell axis vs spatial axis | $\rho = 0.735$; top-100 overlap 52 (chance 5.5) |
| Scoring schemes | CMap vs the two correlations | Cor–Cor $\rho = 0.950$ (433 of top 500 shared); CMap–Cor $\rho = 0.36$–0.38 |
| *Geometric self-consistency* |  |  |
| Scale-bar calibration | adjacent-spot spacing<br>$= 100\ \mu\text{m}$ | 1 px $= 0.2513\ \mu\text{m}$; full panel 10.9 mm (11 mm format) |

#### The reversal acts on specific domains

Using each domain’s own depth-balanced signature as a query, 26 of the 34 candidates fall in the top 100 for D3 and 21 for D5, against a chance expectation of 1.9; the remaining five domains are at or near chance (Fig. 3g). The compounds therefore act on the ECM/interstitial and AT2 domains rather than on the tissue as a whole.

### Gating of the candidate gene list

The same disease axis was scored against a gene-perturbation library of 20,345 perturbations (15) using the authors’ causal predictor. The gating framework applied to the resulting ranking is described here because it converts an ungoverned list into a short one and because it exposes a failure mode that a naive ranking conceals.

A gene-perturbation score is informative only if the effect depends on the cell type being perturbed. Every perturbation was therefore scored on a K562 control axis, and those whose effect was as strong there as on the target axis were removed. Of 9,878 scored entries, 20 were flagged as generalising, 48 were specific to the non-K562 reference, 1,898 were neutral and 7,912 lacked K562 data and are reported as untested. Applying this gate together with the survival-direction filter leaves 25 entries supported by all four scoring routes, 111 by three and 7 by two.

To test whether the candidates act on a compartment rather than on the tissue as a whole, the disease axis was rebuilt for each of 31 subtypes separately and every perturbation re-ranked within each axis. Of 1,466 perturbation entries, counted as gene and direction, reaching the top 100 in at least one cell-type axis, only two reach it in twenty or more of the 31. One is *SFTPC* knockdown, which ranks in the top 100 in 22 of 31 axes but fails the direction check, since lowering a transcript whose high expression is protective does not describe a therapeutic direction. The other is *STAT1* knockdown (21 of 31 axes; hazard ratio 1.08, consistent with the required direction). The remaining candidates are largely axis-specific. The headline number of the ungated ranking is misleading for this reason: ubiquity across cell types is not evidence of a target, and the entry that appears most broadly reproducible does not survive the direction filter.

### 2.8 Analyses that did not work

Six further negative results are reported, each with the reason it is believed to have failed.

#### Gene-level prediction on the pooled axis

Applied to the tissue-pooled disease axis, the pipeline produced a maximum causal score of 0.083 against 1.279 for the official benchmark on the same implementation, and the distribution on our axis has no high tail, whereas the benchmark’s does. Since the method is demonstrably functional, the pooled axis has no detectable match to the expression shifts recorded in the library. The compartment-resolved query does rank, but its survivor count is one. A separate, self-implemented cosine projection, which omits the causal score’s gene-shift term, also ranked genes, but its significance saturates at the permutation floor (237 of 2,439 genes at the minimum attainable *p*) and it cannot discriminate.

#### A developmental trajectory that runs the wrong way

Using the source study’s approach (monocle3 (16) with CytoTRACE (17)), the score is highest rather than lowest in invasive disease: medians are Normal 0.288, AAH 0.374, AIS 0.288, MIA 0.277 and IAC 0.453, a non-monotonic profile with IAC highest. The reversal is robust, with 22 of 23 patients showing a positive per-patient slope and a depth-balanced audit finding IAC highest within every depth stratum. The measurement is nevertheless ambiguous in a way that cannot be resolved. CytoTRACE is constructed from the number of detected genes, so a high score in invasive disease is equally consistent with greater transcriptional disorder and with a less differentiated state; two readings fit the same numbers. Because the analysis measures a stage axis rather than a subtype axis, no AIC/KAC or tumour labels being available, it can be interpreted neither as evidence against the source study’s trajectory nor as evidence for a different one.

#### A spectral mechanism that was an artefact of construction

An initial attempt to read the trajectory from the spectrum of an optimal-transport operator (18, 19) was abandoned after inspection. Under the stage partition the coupling matrix is block upper triangular, so transport does not enter the spectrum; in small independent checks with known row sums the eigenvalue union differed by 0.000 *×* 10^0^ and the number of near-zero eigen-values equalled the cell count of the first four stages. The apparent spectral gap was set by a coupling-weight parameter and not by the data. The result was, on inspection, a property of how the matrix was assembled.

#### H&E imaging at this resolution

A pathology-language model (20) was applied to the H&E images to ask whether the morphological sequence could be read directly from the histology. Across twelve independent configurations (six feature arms crossed with two prompt sets) the best correlation was 0.729, and no setting exceeded 0.74. The ceiling is set by the model architecture: PLIP takes a 224-pixel input yielding 49 tokens, so one token spans about 181 *μ*m of tissue, against the finest available colour source at 5.670 *μ*m per pixel. Above that scale the discriminative power did not vary with detail, and finer sources cannot be tested in this cohort because none exists.

For the use this study would make of it, the decisive result is that the model did not resolve the precursor stages. Per-stage means for AAH, AIS and MIA all fell within 0.15 to 0.22 and interleaved with one another, so the three stages that define the sequence are not separated by this model. What increased with image detail was only the contrast between invasive disease and everything else, with the benign end depressed, rather than a gradient across stages.

A search over 1,344 template-and-term combinations identifies what the model is in fact measuring. The highest raw discrimination, an AUC of 0.972, came from the term “invasive adenocarcinoma with desmoplastic stroma”, but that term also correlated most strongly with tissue density (*ρ* = *−* 0.574), and projecting the density direction out of its embedding reduced the AUC to 0.759. A weaker prompt chosen to be density-neutral retained more of its discrimination under the same projection (raw AUC 0.833, *ρ* = 0.129; projected 0.791). Two independent measurements agree: the lesion score correlates at *ρ* = 0.718 with the tissue fraction of a spot and at *ρ* = −0.794 with an empty-slide control; and the model does not merely fail to rank precursor lesions above normal lung, it inverts the comparison, with AAH versus Normal AUCs of 0.28 to 0.32 across prompt sets. Tumour tissue is itself denser than normal lung, so removing density also removes part of the true signal and the projected values are a lower bound rather than a target. The relevant point is that performance here is not separable from how much tissue a spot contains. To establish that the ceiling reflects resolution rather than the choice of image, every section was registered from the high-resolution image into the CytAssist frame (56 of 56 passing; Methods) and the same spots were rescored from four image sources matched by physical size. The finer source did not improve performance: slice-level correlation was 0.698 for the baseline image, 0.228 for CytAssist, 0.540 for CytAssist contrast-matched to the baseline, and 0.215 for CytAssist downsampled to the baseline’s pixel density. Because downsampling and raw CytAssist perform alike, the difference is not one of pixel density but of white balance and contrast between the two products, and matching the contrast recovers most of it. The conclusion is correspondingly narrow. At this image scale the model cannot perform the task this study would require of it, which is to place a lesion on the precursor-to-invasive axis. That is not a statement that histological detail is uninformative in general: testing that would require a finer colour source than this cohort provides.

#### The ligand layer

NicheNet was used with fibroblasts as receivers, the ten most abundant non-fibroblast cell types of the ECM/interstitial domain as senders, and the same data-derived target set. Only five ligands passed the expression and network gates, and none produced a ligand– target link in the output matrix. Their ligand activities were correspondingly flat, the best corrected area under the precision–recall curve being 0.093 against a baseline of zero. Since NicheNet’s own baseline correction was applied, the failure lies in the candidate set rather than in the scoring: with five ligands the sender panel is too thin for the network to resolve anything. Ligand-level inference is the natural next step, and this is the specific reason it was not pursued here.

#### A finer spatial resolution that measured sections

One attempt to ask whether the RCTD composition space supports more than seven domains pooled spots from all 56 sections into a single *k*-means and cross-validated the result. It returned a clean optimum, but the clusters tracked section identity rather than niche. Pooling across sections makes between-section differences the dominant axis of variation, so the bootstrap measured whether clusters mixed across sections are stable and not whether the composition space supports a finer partition. The analysis was superseded by a per-section clustering with the same *K* scan, which is the version reported above.

## 3 Discussion

The central difficulty of this study is that sequencing depth and tissue density are, in this cohort, the same measurement. A domain’s marker stability, the apparent stage-dependence of the spatial programmes, the cohort-level copy-number difference and the prognostic weight of one domain’s signature all vary with depth, and depth varies with the amount of tumour in a spot. Adjusting for depth is consequently not a neutral correction: it removes part of the biological effect together with the technical one. Adjusted estimates are conservative and may be over-corrected; unadjusted estimates are inflated. Where the choice is consequential, both are reported rather than one being nominated as correct.

The coupling also accounts for the behaviour of the pathology model, which is the one result here obtained without any transcriptomic measurement. Its apparent discrimination was largely a measure of how much tissue a spot contained, and on the axis this study needs, the separation of precursor from invasive lesions, it failed outright. A model that reads tissue density will appear to succeed on this disease for as long as density and malignancy coincide, which is exactly the condition that makes the confound difficult to detect.

This also accounts for the narrower scope of some results relative to the source study. Where a claim depends on comparing dense with sparse tissue, which in this disease means comparing invasive with precursor tissue, the design carries the confound in its structure. The only clean comparisons are those that hold depth fixed, either by pairing within a section or by matching across sections. That is why the one copy-number result retained is a within-section comparison, and why the disease-axis queries were constructed by pairing within patients.

### 3.1 What the positive results establish

Four findings survive the checks applied. First, the tissue resolves into seven recurring niche domains, six of which are ordinary lung structures and only one of which is confined to invasive disease. The lymphoid-aggregate domain is present in precursor stages, which argues against attributing immune aggregation to the onset of malignancy. Second, the ECM/interstitial programme that distinguishes the invasive domain is itself elevated in invasive disease within patients (20 of 22), and the result holds when the genes added by hand are removed and under every leave-one-out variant, so the domain label does not rest on a hand-picked panel. Third, the spatial copy-number signal lies in the AT2 compartment and is present before invasion, consistent with AT2 as the cell of origin and with an orthogonal expression-based classifier. Fourth, a query signature derived from a patient-internal disease axis and filtered on independent survival evidence returns a reproducible set of 34 compounds, dominated by PI3K/mTOR inhibitors, which act on the ECM/interstitial and AT2 domains rather than on the tissue as a whole.

The candidate list should be read as a hypothesis about a compartment rather than as a therapy. Several caveats are structural. The compounds act in cancer cell lines and have not been tested in lung fibroblasts or in any primary tissue. The disease axis against which they were selected is a tissue-level average, so the candidates reverse the aggregate state and need not act on any single cell type. The 34 compounds are the intersection of three scoring schemes, but those schemes share a query and a library and their agreement is therefore a robustness check rather than three independent measurements. The split-half reproducibility of 0.962 describes the stability of the ranking under resampling of patients and says nothing about whether the ranking is biologically correct. No compound reported here has been tested experimentally.

### 3.2 Methodological observations

Several of the checks applied here are reusable, and the difficulties encountered carry lessons that are not specific to this dataset.

A positive control with a known answer, run through the identical pipeline, distinguishes “the method found nothing” from “the data contain nothing detectable by this method”. Without such a control the negative results reported above would be uninterpretable. A related practice is to inspect the distribution of a summary statistic rather than the summary alone: an AUROC of exactly 0.5000 in this study was a degeneracy, all cells carrying one identical value, and not a true null.

Control for a nuisance variable is better served by sensitivity analysis than by a single adjusted estimate. When a measurement is strongly coupled to such a variable, examining subsets of the measurement is more informative than adjusting once. In this study, showing that a result held when the eight hand-added genes of a marker panel were removed, and across all twenty-two leave-one-out variants, established that the finding did not depend on the panel’s arbitrary members. A related caution applies to sub-clustering: because clusters are selected on the same data used to test them, per-subtype contrasts carry a selective-inference penalty that is not corrected here (21), which is one reason the per-compartment gene list is reported as descriptive.

Two further observations concern how criteria are constructed and reported. A gate that every configuration passes is not a gate. The spatial-coherence criterion applied here, which required a partition’s coherence to exceed a label-permutation null on at least 90% of sections, was passed by essentially every configuration tried, including one that uses no spatial information at all (*λ* = 0; coherence 0.54–0.60 against a null 95th percentile of 0.13–0.18). Tissue expression carries spatial autocorrelation intrinsically, so a purely expression-based partition clears the null by a wide margin. The criterion can exclude a grossly anti-spatial partition but cannot demonstrate that spatial information was used; ranking was accordingly decided by cross-seed stability instead. In addition, a criterion introduced by the analysts must not be presented as though it carried external authority. The coherence gate is ours. The source study sets no spatial gate, and the BANKSY authors’ recommendations (3) concern parameters (*λ, k*_geom_, resolution) rather than validation criteria.

Finally, a number produced by an optimiser’s trajectory is not a measurement. One weighting parameter carried for a time in this work, *w*_min_ = (1*/*6) *×* 0.7^*k*^, is the path the solver took rather than a quantity estimated from the data, and should not be reported as though a fit had produced it.

### Relation to the source study

This study set out to extend a published description of the disease sequence (22). The results agree with it on the identity of the niche structures and on AT2 as the compartment carrying copy-number signal, and differ in emphasis on three points. The source study reports a pro-inflammatory niche that is enriched early and less frequent in invasive disease; our spatial arm does not reproduce that direction. Its developmental trajectory places the least differentiated state in precursor lesions, whereas ours runs the other way, although, as noted, the measurement is ambiguous and is not treated as a refutation. The source study also does not describe ambient-RNA correction, whereas one of its samples is, in our hands, dominated by ambient signal. These divergences are reported rather than adjudicated, since each is confounded by a design difference: different platforms, a stage axis rather than a subtype axis, and different approaches to quality control.

### 3.4 Priorities for further work

The comparison that would remove the dominant confound is within a single section, comparing the invasive front with regions of the same section distant from it and holding depth fixed by construction. That analysis was not performed and is the one we would prioritise next, together with testing the candidate compounds in primary lung fibroblasts or in organoids derived from the same patients.

## 4 Limitations

### 1. Depth and density are not separable

Every analysis in this paper is affected by the fact that sequencing depth proxies tissue density. Where the choice is consequential we report both adjusted and unadjusted estimates rather than selecting one.

### 2. No per-cell malignant label

We do not assign malignant status to any cell, spot or domain. The copy-number score is reported as a continuous quantity throughout, and no downstream statistic multiplies it into any other measurement.

### 3. Domain identity is method-sensitive

BANKSY, RCTD composition and marker-based clustering resolve 7, 6 and 5 domains respectively, with an adjusted Rand index of 0.352 against an RCTD-aligned partition. The seven domains are a working description, not a unique decomposition.

### 4. The deconvolution reference bounds what deconvolution can report

The 39-subtype reference has no exhaustion programme, so exhausted T cells are reported under a naive label. T-cell composition in this paper is a composition in the reference’s vocabulary; functional states not present in the reference are invisible to it, and were measured separately by signature projection.

### 5. The *K*^*∗*^ = 7 partition failed its pre-specified stability gate

Across seedings the cross-seed adjusted Rand index never reached 0.90. We report the partition as exploratory and do not use it as a primary result; the consequences are confined to descriptions of domain composition. The companion spatial-coherence criterion we also applied is passed by every configuration tested, including one using no spatial information, and therefore carries no weight in this cohort.

### 6. One clinical stage is represented by one section

The Normal arm of the spatial cohort consists of a single Visium section; MIA is represented by four. Estimates for these stages should not be quoted on their own. Nuclei are more balanced (92,337 normal nuclei across 23 patients), so the imbalance is specific to the spatial arm.

### 7. Two domain identities were revised during the study

The D3 domain was first described as an immune infiltrate, then as a hypoxic/invasive programme, and finally as ECM/interstitial-enriched after depth matching showed the first two readings to be driven by depth. Earlier names for this domain should not be used.

### 8. The marker panel for one analysis was hand-curated

A 22-gene ECM panel used in the early spatial analyses overlaps the depth-balanced marker set by only 14 genes; the remaining eight were added by us. A sensitivity analysis shows the finding is unchanged when those eight are removed, or when any single gene is removed, but the panel itself is not drawn from a single published source.

### 9. Stage comparisons may be contaminated by sample type

Patient-internal pairing removes between-patient variation but not the systematic difference between a precursor lesion and an invasive carcinoma, which is present in every paired comparison. The consistency of a signal across 22 patients does not protect against this, since every patient contributes the same kind of contrast.

### 10. The prognostic analysis is exploratory

Domain marker signatures were used without a signed pre-registration of the parameterisation, and one domain’s result changes substantially when its signature is replaced by a depth-balanced version. Only the direction and its robustness are reported for that domain.

### 11. The compound candidates are untested

They act in cancer cell lines, have not been assessed in lung fibroblasts or primary tissue, and have no experimental validation. They are candidate perturbations.

### 12. The gene-level candidate list rests on one entry

After the specificity gate, the direction filter and the compartment decomposition, only two perturbations rank in the top 100 across twenty or more of the 31 subtype axes, and one of the two fails the direction check. The surviving gene-level claim therefore rests on a single perturbation (*STAT1* knockdown), which we report as an observation rather than as a target.

### 13. The ligand layer produced nothing

NicheNet passed only five ligands, none of which yielded a ligand–target link, so no ligand-level conclusion can be drawn from this cohort. The failure is traceable to a thin sender panel and is not evidence that no ligand effect exists.

### 14. The perturbation libraries are not independent

The gene-level and compound-level libraries share the query signature, and the three compound scoring schemes share both the query and the library. Agreement between them is a robustness check, not independent corroboration. The two libraries are reported side by side and are not used to validate one another.

### 15. The pooled gene-perturbation query produced no signal

Its positive control passes, so the null reflects the query rather than the method, but we cannot distinguish between “this cohort’s pooled disease axis is not represented in the library” and “no single-gene perturbation of this magnitude exists in the library”. Splitting the query by compartment recovers a ranking, so the null is specific to the pooled parameterisation rather than to the library.

### 16. The developmental analysis is ambiguous by construction

The score we use is derived from the number of detected genes, so a high value in invasive disease is equally consistent with greater transcriptional disorder. Two readings fit the same numbers and we cannot separate them with the available labels.

### 17. The imaging analysis is bounded by resolution, not by biology

The H&E-based classifier ceiling is set by the model architecture against the available image scale, and the model does not separate the precursor stages from one another at all. The negative result concerns what this pipeline can extract at this resolution, not what is present in the tissue; a finer colour source would be needed to test the latter.

### 18. No orthogonal validation

No result in this paper has been reproduced in an independent cohort, with a different perturbation library, or experimentally.

## 5 Methods

### 5.1 Data

#### Primary cohort

Two modalities from the same patient series were used. Single-nucleus RNA sequencing (GSE308103) was generated with a fixed RNA-profiling probe panel rather than a standard 3^*′*^ or 5^*′*^ chemistry, so it quantifies a defined set of 18,082 genes rather than the whole transcriptome; this is why housekeeping and mitochondrial transcripts such as *GAPDH*, the ribosomal protein genes and *MALAT1* are absent by design and are not used in any signature. The series contains 75 samples from 25 patients; 798,100 nuclei were called, of which 767,839 passed quality control and 118,894 (15.48%) were called as doublets, leaving **413**,**697 nuclei** profiled over 18,069 genes for analysis. Spatial transcriptomics (GSE307534) was generated on the Visium CytAssist platform in the 11 mm format, 55 *μ*m spots on a 128 *×* 112 grid with a nominal spot-to-spot distance of 100 *μ*m, giving 14,336 spots per section across **56 sections from 25 patients**; the platform implements the spatial-transcriptomics method of Ståhl et al. (23). The two modalities share **23 patients**. The clinical sequence is Normal *→* AAH *→* AIS *→*MIA *→*IAC; the spatial annotations use the source study’s token `LUAD` for the invasive stage, which we map to IAC throughout. Section counts per stage are Normal 1, AAH 11, AIS 14, MIA 4, IAC 26 (Table 5).

**Table 5:** Cohort and data sources. Both modalities are public, come from the same patient series, and share 23 patients. Both are probe-based assays rather than whole-transcriptome chemistries, so each has a defined feature space. The invasive stage is labelled LUAD in the source annotations and is mapped to IAC throughout. Normal is represented by a single spatial section and MIA by four, so estimates for these two stages should not be quoted on their own.

| Item | Single-nucleus RNA-seq | Spatial transcriptomics |
| --- | --- | --- |
| Accession | GSE308103 | GSE307534 |
| Platform | fixed RNA profiling | Visium CytAssist, 11 mm |
| Feature space | 18,069 genes analysed | 18,085 probes |
| Samples | 75 | 56 sections |
| Patients | 25 | 25 |
| Units called | 798,100 nuclei | 14,336 spots per section |
| Retained after QC | 767,839 (96.2%) | 95.03% of spots |
| Doublet rate | 15.48% | — |
| Analysed | **413,697 nuclei** | all 56 sections |
| Grid and spacing | — | 128 × 112; 100 μm |
| Patients in both modalities | 23 |  |

Sections per stage: Normal 1 AAH 11 AIS 14 MIA 4 IAC 26 (source token LUAD)

#### External statistics

Prognostic filtering used TCGALUAD (1) in its archived form: expression and clinical matrices only, giving **483 patients and 177 deaths** (36.6%; 381 patients and 143 events for disease-free survival). The mutation archive in that distribution is a zero-byte file, so no mutation-derived quantity (TMB, driver status) enters any analysis.

#### Perturbation resources

Two libraries were queried. The compound library is LINCS L1000 Level 5 (GSE70138; 118,050 signatures *×* 12,328 genes, 1,826 unique perturbagens across 41 cell lines) (6); we rebuilt it locally as described below rather than querying the CLUE service. The gene-level library is SCMG (15) (20,345 perturbations *×* 18,108 genes), used with the authors’ `CausalGenePredictor` and their reference cell-state manifold.

#### Annotation references

Cell-type annotation was cross-checked against the CellTypist `Human_Lung_Atlas` model (24), itself built on the HLCA (25). Ligand analysis used the NicheNet human ligand–receptor network and ligand–target matrix (26).

#### Resources evaluated and not used

Three external resources were assessed and rejected, and we record them so that their absence is not read as an oversight. An external normal-lung Visium series (GSE248082) was examined as a benign anchor for the copy-number arm; its two usable sections failed a two-way health check (the heldout section scored above threshold in both directions), so no external anchor is claimed. An external lymph-node-metastasis series initially earmarked for the spatial cohort proved, on inspection, to be breast rather than lung cancer and was removed. Three public single-cell cohorts considered in the study’s first design were moved out of scope when the paired series became available, and were never analysed. No GWAS, eQTL, docking or structural data entered this work.

### 5.2 Single-cell processing

After quality control and doublet removal 413,697 nuclei were retained. Contaminant and low-quality clusters were then excluded, leaving 399,579 nuclei in the six mutually exclusive lineage subsets that the atlas is built on. Expression was normalised to counts per 10,000 with a log(1+*x*) transform (CP10K+log1p) throughout. The atlas was built with Seurat (27) using the regularised negative-binomial variance-stabilising transform (28) with `glmGamPoi` (29) for parameter fitting, and integrated across samples with Harmony (30). Doublets were called with scDblFinder (31). An independent second caller was attempted but could not serve as a cross-check: Scrublet (32) reached a maximum score of 0.495 against its automatic threshold of 0.556 on these nuclei and therefore called zero doublets in all 75 samples, a consequence of how sparse these profiles are rather than evidence against the scDblFinder calls. The atlas was clustered per lineage, yielding 39 second-level (L2) subtypes. Endothelial and myeloid marker panels were rebuilt before downstream analysis: the published table provided six canonical genes per endothelial sub-type, which we expanded to as many as twenty genes per subtype across nine endothelial types; the myeloid panel was expanded analogously, drawing on the human lung atlases (25, 33–35) and, for the epithelial subtypes, on the lung adenocarcinoma atlas (2, 36). Automated annotation against the HLCA reference used CellTypist (24). All reported subtype compositions use the expanded panels.

Ambient RNA was assessed in one precursor sample (P24) by three independent checks (patient-level rank of the suspect programme, comparison against a matched negative-control gene set, and an independent readout from a separate region of the same block). The sample was retained with an annotation and all B/plasma-dependent conclusions are reported with and without it.

### 5.3 Spatial domain identification

Spatial domains were identified with BANKSY (3) on each section independently. The main configuration was fixed at *λ* = 0.2, *k*_geom_ = 18 and resolution 0.5, chosen because the authors recommend *λ* ≈ 0.2 for domain segmentation on 55 *μ*m Visium and because higher *λ* values, which the source study used, down-weight the cell’s own expression towards zero. Cross-seed consensus over five seeds produced 434 domain-level clusters across 56 sections; these were collapsed to archetypes by profile-based clustering, giving *K*^*∗*^ = 7 at a subsampling stability of 0.945 (re-clustering on a random 80% of domains, 100 iterations); the cross-seed adjusted Rand index of the per-section domain calls is a separate quantity and is lower (median 0.782). Stability falls below 0.814 at *K* = 8. A descriptive optimum exists at *K* = 6 (0.966) but was not adopted, since the selection rule took the largest *K* passing the stability threshold. Consensus stability was scored with the proportion-of-ambiguous-clustering statistic (37). A spatial-coherence criterion was also computed: per section, the mean coherence of the partition was required to exceed the 95th percentile of a label-permutation null, with at least 90% of sections passing. This criterion is our own construction, not one recommended by the source study or by the BANKSY authors, and it is not used to select the configuration (see Discussion).

### 5.4 Spatial deconvolution

Spot composition was estimated with RCTD (4) against a 39-subtype reference, using the full-section non-epithelial compartment as the reference for the copy-number arm (the arm using each patient’s own benign anchor covers only 10 of 25 cases). Spots were called epithelial when the epithelial subtype had the highest weight; this criterion was fixed before the cohort run. A spot mask was applied at thresholds of 500 counts, 200 genes and 0.15 mitochondrial fraction, removing 4.97% of spots cohort-wide. This is higher than the 1.05% reported by the source study; the difference is attributable to uneven per-section depth rather than to the rule, and is recorded as a known discrepancy.

Two consequences of the mask are guarded explicitly. Per-section removal rates range from 0.02% to 29.6%, so a section in which a domain occupies little area may simply have had more of itself masked; coverage is therefore reported per section and a small domain area is never read as evidence that the niche is rare. And because the 56 sections come from 25 patients, every cross-section summary is clustered by patient, since two or three sections from one patient are not independent samples.

### 5.5 Depth matching

Two independent estimators were used. In the first, spots were binned into per-slide deciles of nUMI and compared only within bins. In the second, each gene was regressed on log_10_(nUMI) per slide with a cubic term, and the domain-versus-rest ranking was recomputed on the residuals. A gene was counted as surviving if it remained in the domain’s top 150 up-genes. The two estimators agree almost gene-for-gene.

### 5.6 Spatial copy-number inference

Copy-number scores were computed with fastCNV (5). The engine discards chrY and chrM, and its classification output is a direction call rather than a malignancy determination; we use only the continuous per-spot score. Domain-level comparisons pair each spot’s score against the median of the remaining spots in the same section, which removes depth by construction. Significance was assessed per domain by a two-sided sign test over sections, with Benjamini–Hochberg control where families of tests are reported (38). Spatial neighbourhood statistics used Squidpy (39). Three further copy-number tools were evaluated and excluded because each requires aligned reads rather than an expression matrix (Numbat (40), HoneyBADGER (41) and CaSpER (42)), as was SCEVAN (43), which was pre-registered but not run.

### 5.7 Prognostic analysis

Survival analysis used TCGA-LUAD (1) (483 patients, 177 deaths; disease-free survival: 381 patients, 143 events, censored at last follow-up), with Cox proportional-hazards models fitted in R’s `survival` package (44). For each domain, the top-30 marker genes were standardised across samples and averaged into a signature score. Univariate Cox models were fitted both continuously (hazard ratio per standard deviation) and on a median split (reference group: low). Hazard ratios for the adjusted models controlled for stage, age and sex.

### 5.8 Query signature construction

For each patient, the disease axis was computed as the mean expression of invasive cells minus that of precursor cells, and the median across patients was taken, giving an 18,069-gene vector, of which 16,104 are represented in the SCMG library. This axis was filtered on independent survival evidence: a gene rising in disease was retained only if its adjusted hazard ratio exceeded 1, and a gene falling in disease only if it fell below 1; all others were set to zero. Of 18,069 genes, 8,799 were retained (51% removed overall; 44% of the 15,792 genes with survival annotation). A second, spatial query axis was constructed by comparing invasive and precursor spots within nUMI deciles per section, merging deciles weighted by the smaller group size, and removing immunoglobulin, mitochondrial and antibody-probe rows.

### 5.9 A local compound-perturbation database

The standard way to query a disease signature against LINCS is the CLUE service (clue.io). Its scoring service was retired on 2026-01-31: the site still serves data downloads, but it no longer accepts scoring jobs, and submission requests are refused (HTTP 401). Rather than depend on a service that can no longer do the work, we rebuilt the query resource locally, which has the further advantage that the exact library state is fixed, versioned and rerunnable rather than whatever the remote service happens to serve on a given day.

The database is a `signatureSearch`-format HDF5 file built directly from the LINCS L1000 Level 5 Zscore matrix (GSE70138, release 2017-03-06). It stores the assay as a genes × signatures array of 12,328 Entrez genes by 118,050 signatures, with columns keyed by perturbagen, cell line and perturbation type; file size is 5.9 GB. Building it through R `(build_custom_db)` was abandoned because the required compression and index handling failed and the per-column writer was prohibitively slow; the file used here is written directly with h5py, which takes 26 s. The database itself is excluded from the repository, being 5.9 GB and regenerable from the two tracked build scripts. The `padj` slot is deliberately omitted because the scoring functions used here do not read it.

Two access paths exist over the same underlying LINCS data, and it matters which analysis used which. The disease-axis query, the spatial-axis query and the splithalf stability check were run through this local database using the official `signatureSearch` package. The domain-signature query and all depth-matched sensitivity runs were run directly on the source matrix through our own implementation of the CMap statistic. Both paths are cross-validated against the official implementation (Table 4).

### 5.10 Slide registration

The CytAssist images are the finer of the two available H&E sources, but Space Ranger ships no transform between them and the spot coordinate frame, so the mapping had to be measured. We built a registration pipeline that places the high-resolution hires image into the CytAssist pixel frame under a similarity transform with an optional horizontal reflection, cytassist = *s* · R(θ) · *M* · hires + *t*.

Three features make the search tractable rather than a blind optimisation. First, the scale is not searched: it is computed analytically for each section from the shipped scalefactors, spred = (fullres μm/px ÷ hires scalefactor) ÷ CytAssist μm/px, with the CytAssist constant 4.6233 μm/px; the search then only refines ±3% around this value, and in the final solution all 56 sections land within 1% of it. Second, the rotation resolves to essentially a binary choice: every section’s fitted θ lies within 5° of 0° or 180°, and the reflection is applied to 53 of 56 sections. Third, the search is a coarse-to-fine pyramid over translation, obtained by phase correlation, scored by a masked normalised cross-correlation of high-pass-filtered greyscale content (search *σ* = 12, verification *σ* = 4).

Registration quality was established by three independent checks rather than one. The adopted transform must score a fine-scale high-pass correlation of at least 0.10 and at least three times the score of the competing reflection branch; this gate separates, by a wide margin, the sections that the initial run failed on (the 52 passing sections have correlation *≥* 0.19 and separation *≥* 7.0, while the four failures had separation *≤* 1.4 and were re-solved with a tissue-outline overlap metric). Second, the same physical spot was cropped from both sources and compared directly, with the coordinate grid propagated through the full transform; the fitted affine reproduces the grid to within 0.18 px everywhere. Third, the residual was measured on the final solution as a translation field: the residual translation is within one pixel on 46 of 56 sections, and three pixels or less on the remainder, and where the field could be fitted it shows no residual rotation (| *ϕ*| *<* 0.1°) and no residual scale (*k* within *±* 0.13%). At the CytAssist scale of 4.6233 *μ*m per pixel this places even the largest residual well inside a spot radius, so no spot is assigned to a neighbouring spot’s tissue. All 56 sections pass. The pipeline was used to align the same spots across image sources for the H&E audit reported in Results; it is retained in the repository.

### 5.11 Perturbation scoring

Compound scoring used the LINCS L1000 Level 5 library (6) (118,050 signatures *×* 12,328 genes; 1,826 compounds) through `signatureSearch` (45) 1.12.0 with three schemes: the CMap weighted-KS statistic (*τ* ; the statistic originates in GSEA (46)) on top-150 up/down gene sets, and Spearman and Pearson correlation on the full numeric vector. Scores were aggregated per compound by the median across cell lines. Gene-level scoring used the SCMG library (20,345 gene perturbations) with the authors’ `CausalGenePredictor` (cos *×* perturbation sign *×* gene-shift *z*), the authors’ reference manifold and their gene standard deviations.

### 5.12 Positive control and numerical verification

The pipeline was verified in two ways before interpretation. Numerically, our weighted-KS implementation was compared against `signatureSearch` 1.12.0 (maximum absolute difference 6.6 *×* 10^*−*14^; signs agreeing for 2000/2000 comparisons; the fast implementation used for the cross-check was `fgsea` (47)), and our vectorised causal score against the authors’ Python implementation (Spearman 1.00000000 over 19,819 entries). Biologically, the official epiblast-to-nascent-mesoderm benchmark was reproduced with the authors’ own manifold and standard deviations; the known regulators rank 1st (TBXT, of 8,450), 2nd (MSGN1), 3rd (MIXL1), 7th (POU5F1), 9th (SNAI1), 11th (EOMES), 12th (NANOG) and 15^th^ (EVX1).

### 5.13 Candidate gating and compartment decomposition

Gene-level candidates passed three additional gates. First, a specificity gate: each perturbation was scored on a K562 control axis, and an entry whose effect there was as strong as on the target axis was flagged as generalising and excluded; entries without K562 data are reported as untested. Second, a compartment decomposition: the disease axis was rebuilt independently for each of 31 subtypes (mean expression difference between invasive and precursor cells within the subtype, median across patients) and every perturbation was re-ranked within each axis; an entry was counted as top-100 for a subtype at that rank. Third, a survival-direction check: a knockdown was kept only for genes whose increased expression predicts worse outcome, and an overexpression only for the converse.

### 5.14 Ligand analysis

Ligand-level analysis used NicheNet (26) with the default human ligand–receptor network and ligand–target matrix. Fibroblasts were the receiver; the sender panel was the ten most abundant non-fibroblast subtypes of the ECM/interstitial domain. The target set was the top-200 up-genes of the data-derived disease axis, not a hand-written list. Candidate ligands were required to be expressed above log(1+CP10K) *>* 0.1 in at least one sender and to be present in the ligand–receptor network.

### 5.15 Reproducibility checks

Split-half reproducibility used four random partitions of patients; each half was scored against LINCS independently, and the Spearman correlation between the two rankings was compared against a null formed by permuting one half’s ranking (300 permutations). Cross-modality agreement compared the single-cell-axis and spatial-axis compound rankings by Spearman correlation and by top-*N* overlap against a chance expectation of *N*^2^*/*118,050. Scale bars in spatial figures were calibrated per section from the median centre-to-centre distance of adjacent in-tissue spots, which is nominally 100 *μ*m; this yields 1 pixel = 0.2513 *μ*m and a full-panel width of 10.9 mm, consistent with the 11 mm capture format.

### 5.16 Software

Analyses used R 4.2.2 (Seurat, spacexr, BANKSY, Squidpy’s R counterparts, `signatureSearch` 1.12.0, survival), Python 3.8 (numpy, scipy, scanpy, h5py) and the SCMG implementation (15, 48). All figure code is in the accompanying repository. Table 6 lists every consequential parameter together with its provenance.

**Table 6:**
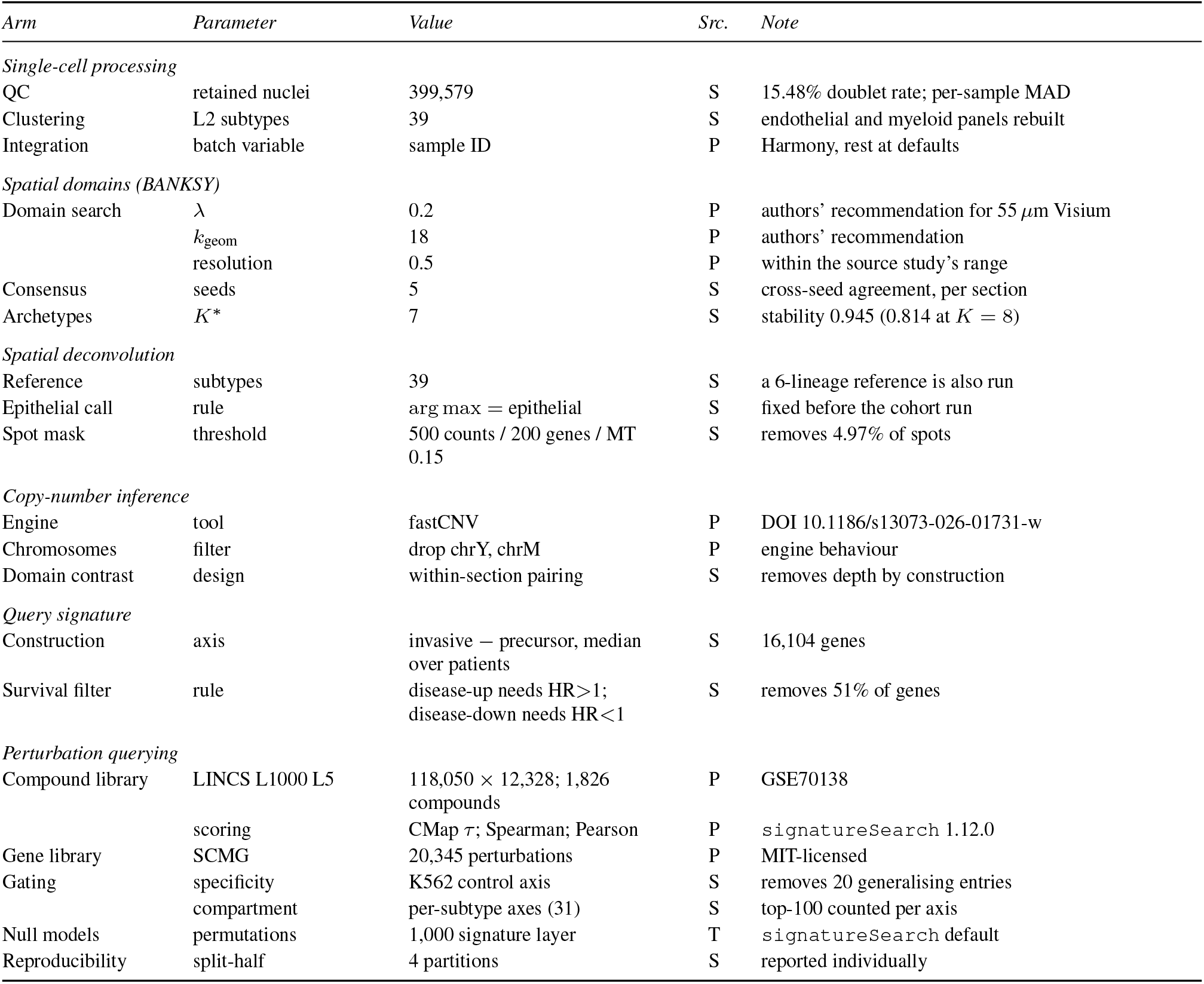
Analysis parameters and their provenance. Each parameter is labelled by its source: **P** = stated in a source paper or software publication, **T** = software default rather than a published recommendation, **S** = fixed by us and recorded in the analysis log. The distinction is drawn because most of the consequential settings in this study are ours rather than the field’s.

## Data and code availability

Raw data are public: single-nucleus RNA sequencing (GSE308103) and Visium spatial transcriptomics (GSE307534). All analysis code, including the scripts that generate every figure, together with the provenance records for each analysis parameter, is available at https://github.com/etonsalmon160-source/luad-invasion-v2. Large intermediate objects are excluded from the repository because they are regenerable from that code; the exclusion list records, for each omitted object, the script that rebuilds it. Supplementary Table S1 lists the 39 subtypes of the deconvolution reference. The other derived tables referred to in the text — per-section copy-number deltas, domain composition summaries and survival model outputs — are provided in the repository alongside the scripts that generate them.

## Acknowledgements

The authors thank the colleagues who supported this work despite the absence of dedicated funding. We are grateful to the senior academic who writes on Bilibili as “Unshakeable Saury”, whose understanding and encouragement sustained the project, and to fellow researchers at Shanghai Jiao Tong University and at Huazhong University of Science and Technology, whose ideas and technical assistance shaped it. We thank the wider academic community for recognising this work and for maintaining the decentralised environment in which it was carried out.

## Author contributions

Z. Li: conceived and designed the study, performed the computational analyses, interpreted the results and wrote the manuscript. **J. Yang:** built and maintained the computing environment, including server resource provisioning, software setup and network configuration, and provided the computational infrastructure on which the analyses were run

## Use of AI tools

Claude Code (Anthropic) and DeepSeek were used to write the analysis code and to polish the language of the manuscript. The study was conceived and designed by the authors; the analytical strategy, the analysis gates and all thresholds were specified by the authors and were not determined by the AI tools. All results passed multi-stage review by the authors before being reported, and the authors take full responsibility for the content of the manuscript.

## Ethics statement

This work is a secondary analysis of de-identified, publicly available human data (GSE308103 and GSE307534). No new human specimens were collected, and no new ethical approval was required. No animals were used.

## Supplementary Table S1

The list follows the reference object used for spatial deconvolution (reference_d.h5 in the accompanying repository). Two entries carry names that require care. *Basophil/Mast 1* is assigned predominantly to epithelial cells, and *Serous* and *Basal* are airway epithelial states rather than the lineages their names might suggest; the lineage-level compositions reported in the main text are computed from the lineage assignment rather than from these names.

**Supplementary Table S1.** The 39 second-level (L2) subtypes of the single-nucleus atlas. Subtypes are listed in descending order of cell number, continued across the two halves of the table. Counts refer to the 399,579 nuclei retained after quality control and are the reference used for spatial deconvolution. These names are not unique identifiers of a lineage–subtype combination: they map onto 67 distinct combinations, and *Basophil/Mast 1* is assigned predominantly to epithelial rather than myeloid cells (main text).

| *Subtype* | *Nuclei* | *Subtype* | *Nuclei* |
| --- | --- | --- | --- |
| AT2 | 88,666 | Vein | 5,654 |
| CD4<sup>+</sup> Naive T | 37,528 | Bronchial Vessel 2 | 4,674 |
| Fibroblast | 35,581 | Goblet/Mucous | 3,587 |
| AT1 | 30,320 | Pericyte | 3,480 |
| Myofibroblast | 24,244 | Lymphatic | 3,354 |
| TREM2<sup>+</sup> Dendritic | 20,571 | CD8<sup>+</sup> Naive T | 2,680 |
| Plasma | 15,975 | EREG<sup>+</sup> Dendritic | 2,476 |
| Macrophage | 10,837 | Basal | 2,386 |
| NK | 10,608 | Nonclassical Monocyte | 2,327 |
| IGSF21<sup>+</sup> Dendritic | 9,193 | Bronchial Vessel 1 | 2,292 |
| Capillary | 8,839 | NKT | 2,172 |
| Vascular Smooth Muscle | 8,408 | Classical Monocyte | 1,985 |
| CD8<sup>+</sup> Mem/Eff T | 8,173 | Myeloid Dendritic Type 1 | 1,953 |
| Capillary Intermediate 1 | 7,194 | CD4<sup>+</sup> Mem/Eff | 1,917 |
| B | 6,940 | Serous | 1,728 |
| Ciliated | 6,697 | Plasmacytoid Dendritic | 925 |
| Artery | 6,388 | Neutrophil | 840 |
| Basophil/Mast 1 | 6,368 | Capillary Aerocyte | 305 |
| Myeloid Dendritic Type 2 | 6,100 | Proliferating Macrophage | 294 |
| Lipofibroblast | 5,920 |  |  |

